# Viewing pictures triggers rapid morphological enlargement in the human visual cortex

**DOI:** 10.1101/408658

**Authors:** K.N.T. Månsson, D.S. Cortes, A. Manzouri, S. Hau, H. Fischer

**Affiliations:** Department of Psychology, Stockholm University, Stockholm, Sweden; Department of Clinical Neuroscience, Karolinska Institutet, Stockholm, Sweden; Department of Psychology, Uppsala University, Uppsala, Sweden

## Abstract

Brain morphology change over the course of weeks, days, and hours, and can be detected by non-invasive structural magnetic resonance imaging. Rapid morphological changes at scanning has yet not been investigated. In a randomized within-group study, high-resolution anatomical images were acquired during passive viewing of pictures or a fixation cross. Forty-seven individuals gray matter volume and cortical thickness were investigated, and both measures increased in the visual cortex while viewing pictures relative to a fixation cross. Thus, brain morphology enlargements were detected in less than 263 seconds. Neuroplasticity is a far more dynamic process than previously shown, suggesting that individuals’ current mental state affects indices of brain morphology. This needs to be taken into account in future morphology studies and in everyday clinical practice.

---

T1-weighted magnetic resonance imaging (MRI) is extensively used in both structural brain imaging research and clinical practice. Non-invasive MRI and T1-weighted images represent a macro-level assessment of brain morphology (*1, 2*), e.g., different tissues, cell types, synapses, and dendritic spines in the cellular milieu. Morphological assessments are routinely used, for example, to assess brain atrophy in normal aging (*3*) and in neurodegenerative disease (*4*). Further, a plethora of human neuroplasticity studies suggest remarkable plastic volumetric alterations induced by motor training (*5*), physical activity (*6*), and by pharmacological agents (*7*). Substantial attention has been directed towards a number of studies on human brain plasticity. For instance, increased gray matter volume in London taxi drivers (*8*) or volumetric changes induced by practicing mindfulness (*9*) or juggling (*10*). Such structural changes in humans are typically found after several weeks, but there is also suggestive evidence of neuroplasticity already after days (*11*) or less. Tost and colleagues (*7*) reported changes after less than two hours, both in striatal gray matter volume and brain response coupling in individuals undergoing an acute dopamine D2 antagonists challenge. Thus, brain morphology is dynamic and alter rapidly by both internal and external environmental influences.

Studies demonstrating changes in less than a few hours raise the question on how fast brain morphology could alter, and if rapid changes at scanning are possible to capture with non-invasive MRI techniques. In contrast to previous studies measuring changes between time points, an alternative approach could be to measure brain morphology alterations more directly and during anatomical scanning. Therefore, to test if the brain’s morphology alters at time of assessment, we conducted a simple experimental manipulation while acquiring T1-weighted anatomical images. In particular, using a randomized balanced within-subject design, 47 healthy participants underwent T1-weighted MRI image acquisition while passively viewing complex arousing pictures or a fixation cross.

## Results

Within-subject voxel-wise whole-brain analysis on voxel-based morphometry (VBM; *12*) gray matter images, found regional and bilateral clusters in the occipital lobe, suggesting that gray matter volume increased in the visual cortex while viewing pictures, see Table 1 and Figure 1 for details. A multi-voxel pattern support vector machine-learning technique also successfully separated VBM images (i.e., picture vs fixation cross), see Supplementary Material for details. A majority of the participants showed an actual increase (above >0%) in left (72.3%, 34/47) and right (68.1%, 32/47) visual cortex gray matter volume. The results remained unchanged when adding global cerebrospinal fluid as a nuisance variable (see also Supplementary Material for other measures on global morphology changes).

**Table 1.** Whole-brain voxel-wise comparisons (paired *t*-test; picture > fixation cross). The analysis included 47 participants.

| Brain regions | MNI |  |  | % Increase | *t* | cluster size (*n* voxels) | *P* <sup>corrected</sup> |
| --- | --- | --- | --- | --- | --- | --- | --- |
|  | *x* | *y* | *z* |  |  |  |  |
| T1-w VBM picture > T1-w VBM fixation cross |  |  |  |  |  |  |  |
| Left visual cortex cluster |  |  |  | 1.4% |  | 705 | <0.020 |
| Lingual gyrus | -18 | -86 | -9 |  | 4.57 |  |  |
| Fusiform gyrus | -24 | -86 | -15 |  | 4.35 |  |  |
| Right visual cortex cluster |  |  |  | 1.3% |  | 789 | <0.020 |
| Lingual gyrus | 12 | -88 | -9 |  | 4.29 |  |  |
| Fusiform gyrus | 33 | -76 | -14 |  | 4.50 |  |  |

**Abbreviations:** T1-w, T1-weighted; VBM, voxel-based morphometry; MNI, Montreal Neurological Institute space

**Fig. 1.**
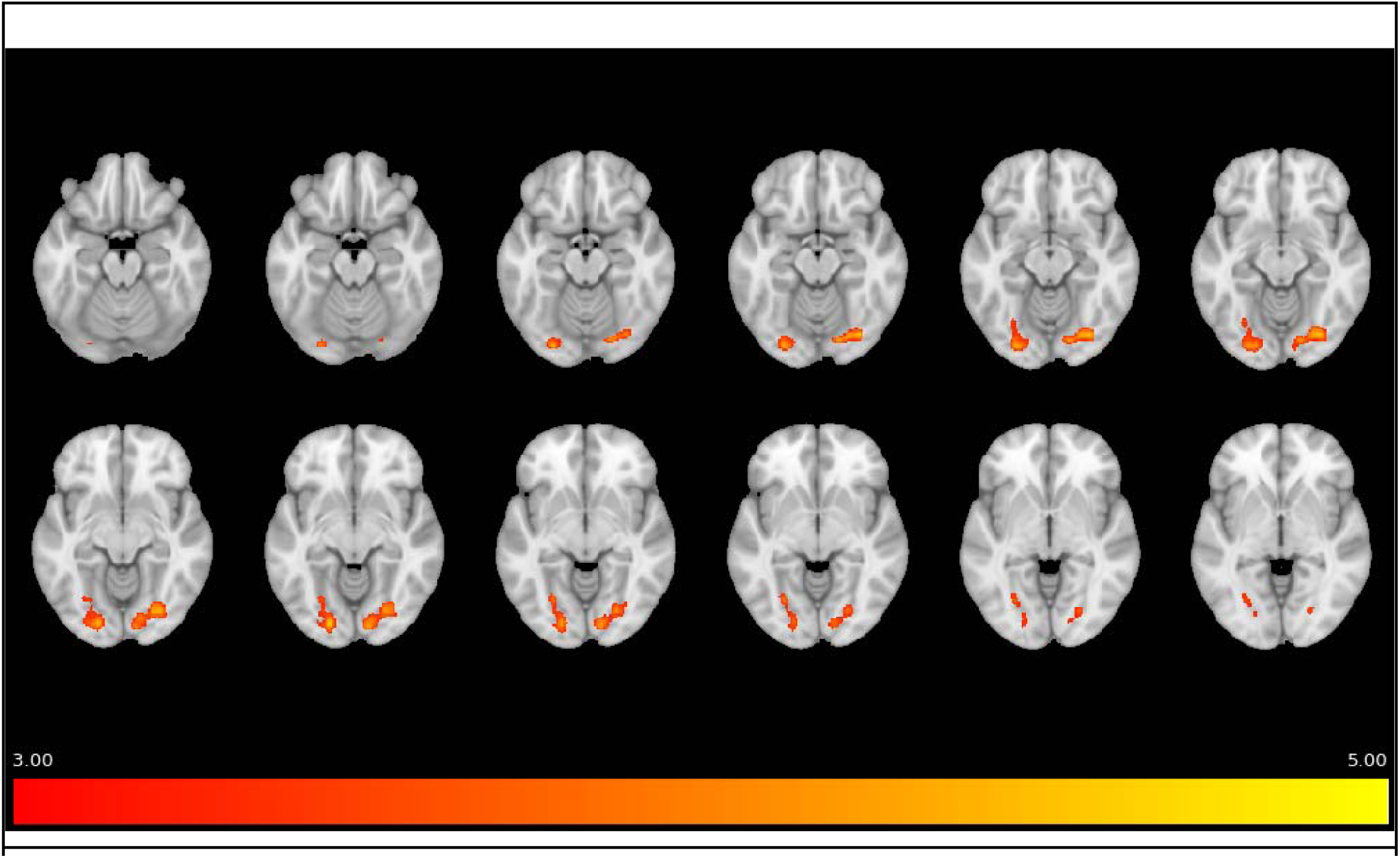
Voxel-based morphometry gray matter volume enlargements while viewing pictures. Whole-brain analysis on voxel-based morphometry gray matter images demonstrating rapid regional changes (≤263 seconds) in bilateral clusters of the occipital lobe, suggesting volumetric increase in the visual cortex while viewing pictures. Whole-brain plot demonstrating significant changes while participants were viewing complex arousing pictures relative to a fixation cross (the figure only include clusters surviving *P*<0.001, *k* threshold >411 voxels). The color bar represents *t*-values ranging from three (red) to five (yellow).

In addition to VBM, by use of a surfaces-based method (implemented in Freesurfer; *13*) to measure cortical thickness, increased occipital lobe thickness was found (see also Supplementary Material).

## Discussion

Forty-seven healthy individuals were assessed while being scanned with MRI under two conditions (randomized to start with either viewing complex arousing pictures or a fixation cross). The study provides evidence of rapid morphological changes in less than 263 seconds in the human “visual cortex” (i.e., lingual and fusiform gyrus of the occipital cortex) while passive viewing of complex arousing pictures. This was demonstrated using both VBM and a surface-based cortical thickness preprocessing technique, as well as univariate and multivariate analytics.

Previous studies that used noninvasive neuroimaging in humans and non-human primates have suggested gray matter volume changes within hours (the T1-weighted anatomical measurements were separated by approximately two hours). For example, seven individuals had pharmacotherapy with the dopamine D2 antagonist haloperidol (*7*), and ten rhesus monkeys were exposed to antiepileptic drugs (*14*). Further, enlargement in the visual cortex gray matter have been shown after less than two hours of training in a color naming learning task, still, the brain measurements were separated by three days (*11*). Evidence from these studies indicate that brain morphology alters in response to both internal and external environmental influences, within days or hours of exposure. We add that morphological changes occur and can be detected while being scanned.

If the current findings represent dynamics in the brain’s gray matter volume and cortical thickness, what specifics in morphology change that fast? Animal research offers some insights on rapid brain morphology change and dendritic spine plasticity is one likely target, for a review see Alvarez and Sabatini (*15*). For example, visual cortex spine plasticity in response to altered sensory inputs has been demonstrated in an adult mouse model (*16*). Also, recent techniques in optogenetics have found evidence of dendritic spine morphology enlargements in less than two seconds (*17*). However, it still remains unclear what brain volume/thickness derived from T1- weighted images truly reflects, for a review see Tardif et al. (*1*). Neuroplasticity studies (e.g., *18, 19*) suggest that the macroscopic assessment of gray matter volume with T1-weighted imaging represent a complex mixture of different cellular mechanisms, e.g., plasticity in synapses, neurons, and glial cells (*2*). Another possible explanation is that changes in blood volume lead to local increases in measures of gray matter volume, e.g., brain oxygenation could change the T1- weighted images (*20*). A study of 28 healthy individuals found no change in gray matter volume after ketamine administration relative to placebo (*21*), and the authors stressed the need of randomized controlled trials because volume changes could be attributed to general changes in physiology such as blood flow. Further, Franklin et al. (*22*) exposed 15 individuals to a single-dose of the antispastic drug baclofen and found a partial overlap between pre-post changes (within about two hours) in gray matter volume and cerebral blood flow. The authors suggested that change in blood flow is masquerading as reductions in gray matter volume. Yet, there are some limitations of this previous research. To determine commonalities between two types of assessments it would be crucial to know the specific effects of such drug on brain function and volume separately. Importantly, an overlap between these two measures is not necessarily evidence that they capture the same signal. In the current study, change in visual cortex morphology occur while the individual processed visual stimuli, thus reflecting a direct assessment of volumetric change in key brain regions involved in visual stimuli processing. It is possible that the current findings also can be related to increased blood flow, and if so, this study provides a rationale to perform more direct assessments of brain morphology while the participants engage in functional tasks. The sensitivity of morphology observed in our study suggests that T1-weighted imaging could be regarded as a hybrid measure of both brain function and brain morphology. This paradigm change could inform novel structural-functional image acquisition sequences for use in future studies and clinical practice.

However, rapid change in morphology can only to some degree be sensitive to the brain’s current actions or blood flow, otherwise T1-weighted derived brain anatomy would not yield such accurate predictions of chronological age (i.e., ±5 years) (*23, 24*). Hence, brain morphology appears to be both dynamic and stable over time. The rapid dynamic feature of volume and thickness, as found in the current study, also suggests that T1-weighted images, to some degree, may be affected by the individual’s current mental state (e.g., thoughts and feelings). Because all participants underwent two image acquisitions with visual conditions of different complexity, and the order of presentation was randomized across participants, carry-over effects from one condition to the other are not likely. The current study design and results suggest that the morphological change was bidirectional, i.e., both volume/thickness increases (fixation<pictures) and reductions (pictures>fixation). If brain volume and thickness are affected by mental state it is possible that previous experimental tasks can affect the measure of brain morphology, similar to what has been shown to impact resting-state BOLD-fMRI (*25*). However, this research question goes beyond the current study. Nonetheless, it is common to offer participants music, radio, or even movies while acquiring anatomical images. Such conditions could potentially impact measures of gray matter volume and cortical thickness. This is certainly an issue that also needs to be considered in multi-site studies, e.g., consortiums with structural brain images where the procedure for acquiring the MRI data may vary across sites.

It is possible that issues related to image registration and head motion could confound the current results. This seems particularly important because participants may have moved more while viewing attention-capturing arousing pictures, as compared to a stable fixation cross. However, we carefully checked all raw structural images and excluded five individuals due to heavy motion, i.e., ringing artefacts or lower quality, as indicated in the image preprocessing steps. Moreover, no difference in image quality between the conditions was found. Reuter and colleagues reported that gray matter volume estimates could be biased by head motion, and showed that motion appears as loss of volume (*26*). In contrast, the current study only found enlargement, and not reductions, of occipital gray matter volume while viewing pictures. Further, effects related to motion are expected to appear by the edges of brain tissue, which was not evident in the current study.

This study was randomized and counterbalanced, all individuals were assessed at multiple time points using a within-subject design and stimuli selected to maximize the difference in visual complexity. Two independent image preprocessing pipelines and statistical models point in the same direction. The output of the multivariate support vector machine-learning classifier suggests that there is a complex pattern of volume that separates the T1-weighted images, and such algorithm can be used to predict data at the individual level. Moreover, there are multiple techniques to analyze structural T1-weighted images and we found similar effects with VBM and a surface-based technique to assess cortical thickness.

To our knowledge this is the first morphological study in humans to directly assess rapid changes in gray matter volume and cortical thickness during an ordinary T1-weighted scanning session. Importantly, we found that indices of brain morphology changed rapidly in 263 seconds, suggesting that the interpretation of gray matter volume and cortical thickness as morphometric trait measures needs to be revised.

## Acknowledgments

We are grateful to Tie-Quing Li, Natalie Ebner and Tomas Furmark for providing valuable comments on the current manuscript.

## Authors contributions

Funding acquisition: SH and HF. Conceptualization: KNTM, HF and SH. Project administration: DSC and KNTM. Methodology, formal analysis and software: KNTM and AM. KNTM wrote the original draft, and HF, SH, DSC and AM were involved in reviewing and editing. All authors read and approved the final manuscript.

## Data and material availability

All data in the manuscript and the supplementary materials is available upon request.

## List of Supplementary Materials

Supplementary Material and Methods

Supplementary Text

Fig. S1

References (27 - 33)

## References

1. C. L. Tardif et al., Advanced MRI techniques to improve our understanding of experience-induced neuroplasticity. Neuroimage. 131, 55–72 (2016).

2. E. Wenger, C. Brozzoli, U. Lindenberger, M. Lövdén, Expansion and Renormalization of Human Brain Structure During Skill Acquisition. Trends Cogn. Sci. 21, 930–939 (2017).

3. L. Nyberg et al., Longitudinal evidence for diminished frontal cortex function in aging. Proc. Natl. Acad. Sci. U. S. A. 107, 22682–22686 (2010).

4. T. L. S. Benzinger et al., Regional variability of imaging biomarkers in autosomal dominant Alzheimer’s disease. Proc. Natl. Acad. Sci. U. S. A. 110, E4502–9 (2013).

5. E. Wenger et al., Repeated Structural Imaging Reveals Nonlinear Progression of Experience-Dependent Volume Changes in Human Motor Cortex. Cereb. Cortex. 27, 2911–2925 (2017).

6. K. I. Erickson et al., Exercise training increases size of hippocampus and improves memory. Proc. Natl. Acad. Sci. U. S. A. 108, 3017–3022 (2011).

7. H. Tost et al., Acute D2 receptor blockade induces rapid, reversible remodeling in human cortical-striatal circuits. Nat. Neurosci. 13, 920–922 (2010).

8. E. A. Maguire et al., Navigation-related structural change in the hippocampi of taxi drivers. Proc. Natl. Acad. Sci. U. S. A. 97, 4398–4403 (2000).

9. B. K. Hölzel et al., Mindfulness practice leads to increases in regional brain gray matter density. Psychiatry Res. 191, 36–43 (2011).

10. B. Draganski et al., Neuroplasticity: changes in grey matter induced by training. Nature. 427, 311–312 (2004).

11. V. Kwok et al., Learning new color names produces rapid increase in gray matter in the intact adult human cortex. Proc. Natl. Acad. Sci. U. S. A. 108, 6686–6688 (2011).

12. C. Gaser, R. Dahnke, CAT-a computational anatomy toolbox for the analysis of structural MRI data. HBM. 2016, 336–348 (2016).

13. A. M. Dale, B. Fischl, M. I. Sereno, Cortical surface-based analysis. I. Segmentation and surface reconstruction. Neuroimage. 9, 179–194 (1999).

14. Y. Tang et al., Single-dose intravenous administration of antiepileptic drugs induces rapid and reversible remodeling in the brain: Evidence from a voxel-based morphometry evaluation of valproate and levetiracetam in rhesus monkeys. Neuroscience. 303, 595–603 (2015).

15. V. A. Alvarez, B. L. Sabatini, Anatomical and physiological plasticity of dendritic spines. Annu. Rev. Neurosci. 30, 79–97 (2007).

16. T. Keck et al., Massive restructuring of neuronal circuits during functional reorganization of adult visual cortex. Nat. Neurosci. 11, 1162–1167 (2008).

17. S. Yagishita et al., A critical time window for dopamine actions on the structural plasticity of dendritic spines. Science. 345, 1616–1620 (2014).

18. M. Lövdén, E. Wenger, J. Mårtensson, U. Lindenberger, L. Bäckman, Structural brain plasticity in adult learning and development. Neurosci. Biobehav. Rev. 37, 2296–2310 (2013).

19. E. Dayan, L. G. Cohen, Neuroplasticity subserving motor skill learning. Neuron. 72, 443–454 (2011).

20. B. Haddock, H. B. W. Larsson, A. E. Hansen, E. Rostrup, Measurement of brain oxygenation changes using dynamic T(1)-weighted imaging. Neuroimage. 78, 7–15 (2013).

21. A. Höflich et al., Imaging the neuroplastic effects of ketamine with VBM and the necessity of placebo control. Neuroimage. 147, 198–203 (2017).

22. T. R. Franklin et al., A VBM study demonstrating “apparent” effects of a single dose of medication on T1-weighted MRIs. Brain Struct. Funct. 218, 97–104 (2012).

23. G. Ball, C. Adamson, R. Beare, M. L. Seal, Modelling neuroanatomical variation during childhood and adolescence with neighbourhood-preserving embedding. Sci. Rep. 7, 17796 (2017).

24. K. Franke, G. Ziegler, S. Klöppel, C. Gaser, Alzheimer’s Disease Neuroimaging Initiative, Estimating the age of healthy subjects from T1-weighted MRI scans using kernel methods: exploring the influence of various parameters. Neuroimage. 50, 883–892 (2010).

25. K.-C. Tung et al., Alterations in resting functional connectivity due to recent motor task. Neuroimage. 78, 316–324 (2013).

26. M. Reuter et al., Head motion during MRI acquisition reduces gray matter volume and thickness estimates. Neuroimage. 107, 107–115 (2015).

27. Lang, PJ, International affective picture system (IAPS): affective ratings of pictures and instruction manual. Technical Report A-6 (2005).

28. H. Fischer et al., Brain habituation during repeated exposure to fearful and neutral faces: a functional MRI study. Brain Res. Bull. 59, 387–392 (2003).

29. J. Ashburner, K. J. Friston, Diffeomorphic registration using geodesic shooting and Gauss-Newton optimisation. Neuroimage. 55, 954–967 (2011).

30. M. Reuter, N. J. Schmansky, H. D. Rosas, B. Fischl, Within-subject template estimation for unbiased longitudinal image analysis. Neuroimage. 61, 1402–1418 (2012).

31. J. Schrouff et al., PRoNTo: Pattern recognition for neuroimaging toolbox. Neuroinformatics. 11, 319–337 (2013).

32. S. Klöppel et al., Automatic classification of MR scans in Alzheimer’s disease. Brain. 131, 681–689 (2008).

33. M. Misaki, W.-M. Luh, P. A. Bandettini, The effect of spatial smoothing on fMRI decoding of columnar-level organization with linear support vector machine. J. Neurosci. Methods. 212, 355–361 (2013).

